# It’s okay to be green: Draft genome of the North American bullfrog (Rana [Lithobates] catesbeiana)

**DOI:** 10.1101/100149

**Authors:** S. Austin Hammond, René L. Warren, Benjamin P. Vandervalk, Erdi Kucuk, Hamza Khan, Ewann A. Gibb, Pawan Pandoh, Heather Kirk, Yongjun Zaho, Martin Jones, Andrew J. Mungall, Robin Coope, Stephen Pleasance, Richard A. Moore, Robert A. Holt, Jessica M. Round, Sara Ohora, Branden V. Walle, Nik Veldhoen, Caren C. Helbing, Inanc Birol

## Abstract

Frogs play important ecological roles as sentinels, insect control and food sources. Several species are important model organisms for scientific research to study embryogenesis, development, immune function, and endocrine signaling. The globally-distributed Ranidae (true frogs) are the largest frog family, and have substantial evolutionary distance from the model laboratory *Xenopus* frog species. Consequently, the extensive *Xenopus* genomic resources are of limited utility for Ranids and related frog species. More widely applicable amphibian genomic data is urgently needed as more than two-thirds of known species are currently threatened or are undergoing population declines.

Herein, we report on the first genome sequence of a Ranid species, an adult male North American bullfrog (*Rana [Lithobates] catesbeiana*). We assembled high-depth Illumina reads (66-fold coverage), into a 5.8 Gbp (NG50 = 57.7 kbp) draft genome using ABySS v1.9.0. The assembly was scaffolded with LINKS and RAILS using pseudo-long-reads from targeted *denovo* assembler Kollector and Illumina Synthetic Long-Reads, as well as reads from long fragment (MPET) libraries. We predicted over 22,000 protein-coding genes using the MAKER2 pipeline and identified the genomic loci of 6,227 candidate long noncoding RNAs (lncRNAs) from a composite reference bullfrog transcriptome. Mitochondrial sequence analysis supported *Lithobates* as a subgenus of *Rana.* RNA-Seq experiments identified ~6,000 thyroid hormone– responsive transcripts in the back skin of premetamorphic tadpoles; the majority of which regulate DNA/RNA processing. Moreover, 1/6^th^ of differentially-expressed transcripts were putative lncRNAs. Our draft bullfrog genome will serve as a useful resource for the amphibian research community.

## Introduction

Living in the most varied environments with both aquatic and terrestrial life stages, frogs are known to be evolutionary innovators in responding to challenges. However, diseases and infections such as chytrid fungus (Raffel et al. 2013), iridovirus (Lesbarrères et al. 2012), and trematode parasites (Hayes et al. 2010) are causing local and regional die-offs. In tandem with habitat loss, which is exacerbated by climate change, these factors have resulted in a worldwide amphibian extinction event unprecedented in recorded history: over two-thirds of ~7,000 extant species are currently threatened or declining in numbers (AmphibiaWeb 2017).

Frogs are important keystone vertebrates, and have provided key discoveries in the fields of ecology, evolution, biochemistry, physiology, endocrinology, and toxicology (Helbing 2012). Yet, there are huge data gaps for understanding their basic biology at the molecular level; few frog genomes are available, and none represent a member of the Ranidae (true frogs), the largest frog family with species found on every continent except Antarctica. The North American bullfrog, *Rana (Lithobates) catesbeiana*, is an ideal species for building a representative Ranid genomic resource because it is consistently diploid, and has the widest global distribution of any true frog. Originally from eastern North America, the bullfrog has been introduced throughout the rest of North America, South America, Europe and Asia. It is farmed for food in many locations worldwide, and is considered an invasive species in several regions (Liu and Li 2009).

The genomes of two *Xenopus* species (*X. tropicalis* and *X. laevis*) have been sequenced and annotated to some extent (Buisine et al. 2015; Session et al. 2016), but these Pipids have an estimated divergence from the Ranidae ~260 million years ago (MYA) (Sumida et al. 2004). This evolutionary separation is accentuated by their differing life histories, behavior, markedly different sex differentiation systems (Eggert 2004); and recent evidence suggests that the innate immune system of *Xenopus* is fundamentally different from three frog families including the Ranidae (Kiemnec-Tyburczy et al. 2012). As a consequence, the degree of sequence is variation is such that *Xenopus* genomic and transcriptomic resources are inadequate for studying Ranid species (Helbing 2012). The genome of a more-closely related frog, the Tibetan Plateau frog (*Nanorana parkeri*), has been recently released (Sun et al. 2015), though this species is also substantially separated from Ranids by approximately 89 million years (timetree.org).

Using some of the latest sequencing and bioinformatics technologies, we have sequenced, assembled, and annotated an initial draft sequence of the ~5.8 billion nucleotide North American bullfrog genome (NG50 length 57,718 bp). We predicted 52,751 transcripts from 42,387 genes, of which 22,204 had supporting biological evidence, and were deemed high confidence. We anticipate that this much-needed resource, which we make public alongside comprehensive transcriptome assembly data, will directly and immediately impact genetic, epigenetic, and transcriptomic bullfrog studies. On a wider scale, it will enable developmental biology research ranging from amphibians to mammals, provide direly needed insights to curb rapidly declining Ranid populations, and further our understanding of frog evolution.

## Results

The draft assembly of the *R. catesbeiana* genome consists of 5.8 Gbp of resolved sequence (Table 1). The majority of raw reads from the PET libraries were successfully merged (63–75%), yielding longer pseudo-reads (mean +/− SD, 446 +/−107 bp). The success of the pre-assembly read merging allowed us to use a value of the assembly k parameter greater than our shortest PET read length, increasing our ability to resolve short repetitive sequences. Genome scaffolding with orthogonal data, which included the reference transcriptome and scaffolds assembled at a lower k value, greatly improved the contiguity of the resulting assembly (Table 1). We assessed the improvement to the assembly after each round of scaffolding using the NG50 length metric and the number of complete and partial core eukaryotic genes (CEGs) using CEGMA, which reports a proxy metric for assembly completeness in the genic space (Parra et al. 2009). Using the Synthetic Long-Reads (SLR) and the Kollector (Kucuk, et al., *in press*) targeted gene assembly (TGA) tool, RAILS (Warren 2016) merged over 56 thousand scaffolds; this permitted the recovery of an additional four partial CEGs, and raised the contiguity of the assembly to approximately 30 kbp (Supplemental Table S1). The most dramatic improvements to assembly contiguity and resolved CEGs were obtained using LINKS (Warren et al. 2015a) and the MPET reads (NG50 increase of ~16 kbp and 10 additional complete CEGs; Supplemental Table S1), followed by the combined Kollector TGA and the lower-k whole genome assembly (~8 kbp improvement to NG50 and 9 additional complete CEGs; Supplemental Table S1).

**Table 1.** Assembly statistics for sequences 500 bp or more in length

|  | Unitig | Contig | Scaffold |
| --- | --- | --- | --- |
| Number ≥ 500 bp | 2,737,303 | 2,191,947 | 1,548,590 |
| Number ≥ N50 | 420,964 | 295,271 | 30,262 |
| Number ≥ NG50 | 438,623 | 300,168 | 22,767 |
| N80 (bp) | 1,252 | 1,862 | 2,934 |
| N50 (bp) | 3,620 | 5,302 | 44,121 |
| NG50 (bp) | 3,509 | 5,239 | 57,718 |
| N20 (bp) | 8,198 | 11,740 | 155,955 |
| Max (bp) | 68,999 | 90,443 | 1,375,273 |
| Reconstruction (Gbp) | 5.715 | 5.787 | 5.842 |

The automated gap closing tool Sealer (Paulino et al. 2015) was used twice during the assembly process. First, prior to the rounds of rescaffolding to increase the amount of resolved sequence available to inform the scaffolding algorithms, and then post-rescaffolding to improve the sequence contiguity and content for the MAKER2 gene prediction pipeline (Holt and Yandell 2011). Sealer closed 55,657 gaps, and resolved nearly 9 Mbp of sequence in the initial scaffolds, and further resolved 20 Mbp of sequence in its second round, closing 61,422 additional gaps.

We identified more than 60% of the *R. catesbeiana* genome as putative interspersed repeats (Supplemental Table S2). The draft bullfrog genome includes 101 (40.7%) “complete” CEGMA genes, and 212 “complete or partial genes” (85.5%). Application of the MAKER2 genome annotation pipeline to the final draft assembly resulted in a set of 42,387 predicted genes and 52,751 transcripts. The criteria applied to identify the high confidence set of genes reduced the population by approximately half, to 22,204 genes and 25,796 transcripts (Supplemental Fig. S1). Of this high confidence set, 15,122 predicted proteins encoded by 12,671 genes could be assigned a functional annotation based on significant similarity to a SwissProt entry.

Furthermore, 680 proteins from 590 genes were identified as particularly robust predictions by GeneValidator (Dragan et al. 2016) (score ≥ 90). This ‘golden’ set includes several members of the Homeobox (HOX), Forkhead box (FOX), and Sry-related HMG box (SOX) gene families, which are transcription factors involved in developmental regulation (Kamachi and Kondoh 2013; Mallo and Alonso 2013; Schmidt et al. 2013). Immune-related genes, including interleukins 8 and 10, interferon gamma, and Toll-like receptors 3 and 4 were also confidently annotated.

### IncRNA

The discovery and analysis of lncRNAs represents a new frontier in molecular genetics, and is of major relevance to the biology behind the functional and largely unexplored component of the transcriptome. The low degree of lncRNA primary sequence conservation between organisms, and the lack of selective pressure to maintain ORF integrity or codon usage complicates traditional similarity based discovery methods (Quinn and Chang 2016). The subtractive approach to lncRNA detection that we employed identified 6,227 candidate lncRNA sequences.

### Differential expression

Characterization of the regulatory factors that mediate thyroid hormone (TH) dependent initiation of tissue specific gene expression programs during metamorphosis have been extensively studied in *X. laevis*. However, this species experiences markedly different environmental conditions in its natural habitat than many Ranids do, and these experiments employed supraphysiological levels of TH (Buckbinder et al. 1992; Wang et al. 1993). Our present analysis of the TH-induced metamorphic gene expression program in the back skin detected nearly 6,000 genes significantly (p < 0.05) differentially expressed upon T3 (the TH 3,3’,5-triiodo-L-thyronine) treatment (Fig. 1), including those found previously through targeted quantitative polymerase chain-reaction (qPCR) experiments (Supplemental Table S3). The most prominent “biological process” gene ontologies associated with the Swiss-Prot derived functional annotations are related to RNA/DNA processing, signal transduction (including hormone signaling), and functions related to cell growth and division (Supplemental Fig. S2). A selection of new transcripts related to RNA/DNA processing were evaluated using qPCR and found to show similar relative abundance as observed with the RNA-Seq data (Supplemental Fig. S3; Supplemental Table S3).

**Figure 1.**
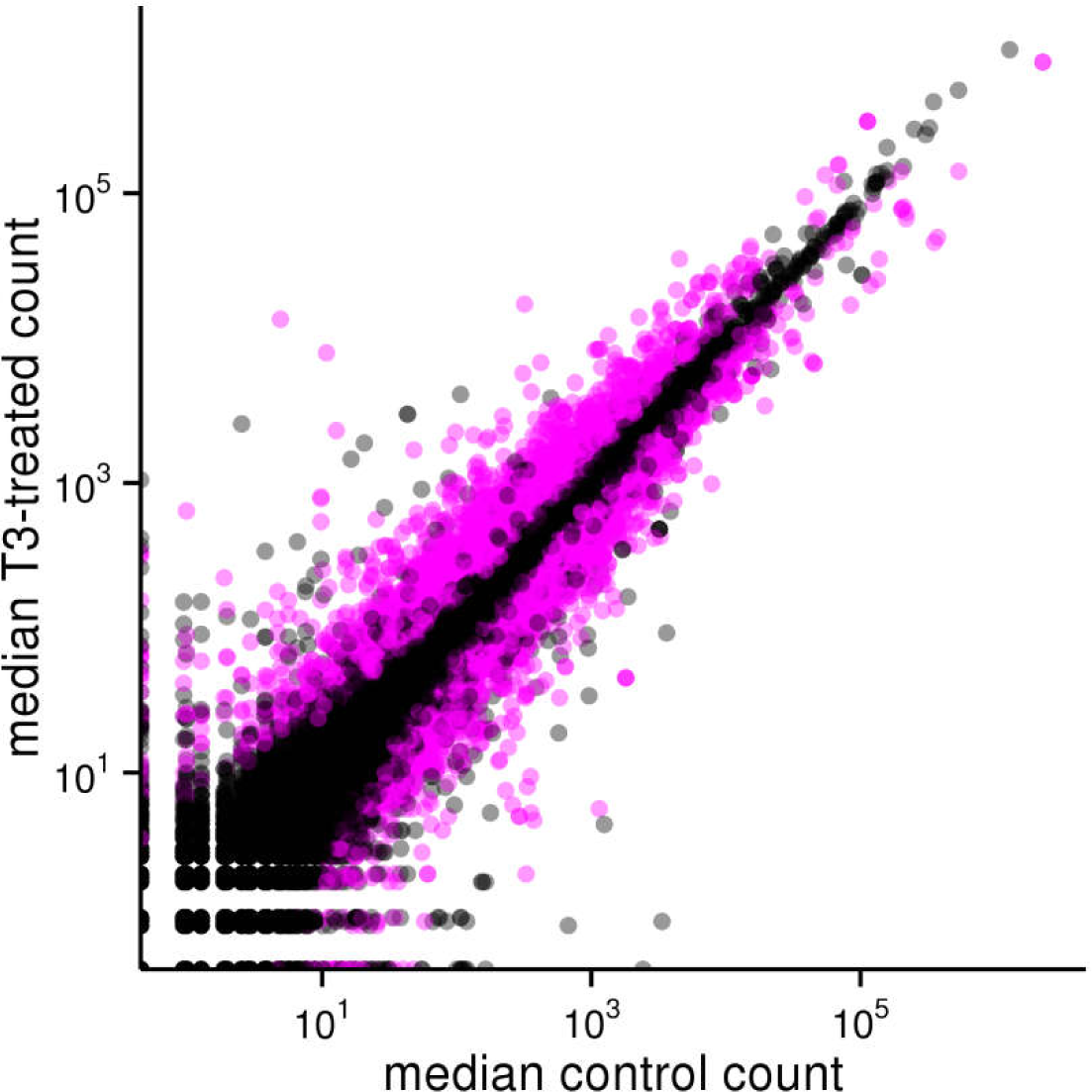
Median counts of genes detected in the back skin of premetamorphic *R. catesbeiana* tadpoles treated with vehicle control or T3 for 48h. Gene transcripts determined to be significantly differentially expressed (DESeq2 adjusted p-value < 0.05) are indicated in pink.

The effect of T3 treatment was not limited to the predicted protein coding genes, as expression of 1,085 candidate lncRNAs was also significantly affected. A selection of lncRNA transcripts was evaluated using qPCR (Supplemental Fig. S4).

### Non-iridovirus sequence in bullfrog

Significant similarity to a frog virus 3 (FV3) protein was noted for one of the gene annotation. FV3 is the type species of the Ranavirus genus, so we explored the possibility that a FV3-like viral genome had wholly or partially integrated into the bullfrog specimen we sequenced. The whole-genome and transcript alignments showed limited similarity between a single bullfrog scaffold and two FV3 sequences that encode hypothetical proteins, FV3gorf5R and FV3gorf98R (not shown). The region of similarity was identified as part of a conserved US22 domain (not shown).

### Phylogenetic analysis of amphibian mitochondrial gene and genomes

Frog taxonomy is subject of debate (Pauly et al. 2009). To address the controversy, we performed a number of phylogenetic experiments comparing selected amphibian mitochondrion (Mt) genomes and Mt genes at the nucleotide level (Fig. 2). As expected, we observe clear separation of salamanders and toads (genus *Bufo*) from other species as outgroups (Fig. 2A; Supplemental Figs. S5-S8). We color-coded the *Lithobates* and *Rana* in yellow and blue, respectively, as re-classified by Frost et al. (2006), to identify the relative genus positioning within the generated phylogenetic trees. At least at the genetic level, the *Lithobates* group branches out of the *Rana* group, as opposed to forming a distinct clade such as what is observed for salamanders and toads. Comparing specific frog Mt gene *cyb*, *Rana* and *Lithobates* often branch together indicative of the close genetic conservation of these species, but do not form independent clades, which suggests a high degree of sequence conservation instead of the divergence observed between distinct genera (Fig. 2B; Supplemental Fig. S6). Ribosomal RNA genes *rnr1* and *rnr2* show phylogenies similar to that of the entire Mt genome, this time with *Rana* branching out of the *Lithobates* clade (Fig. 2C, 2D; Supplemental Figs. S7, S8).

**Figure 2.**
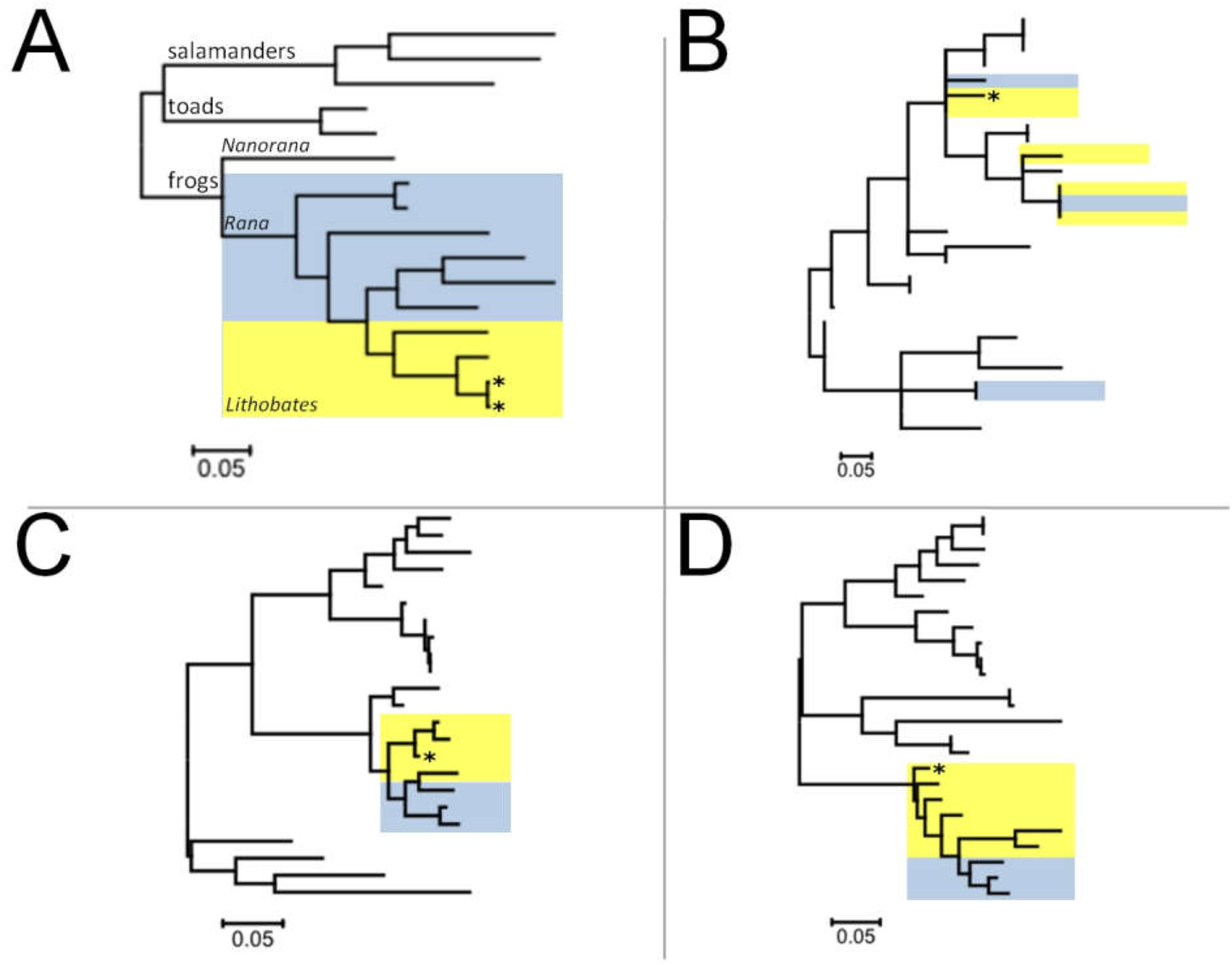
Molecular phylogenetic analysis of amphibian Mitochondrial genomes and genes by Maximum Likelihood method. The evolutionary history was inferred by using the Maximum Likelihood method based on the Tamura-Nei model (Tamura and Nei 1993). The tree with the highest log likelihood is shown. Initial tree(s) for the heuristic search were obtained automatically by applying Neighbor-Join and BioNJ algorithms to a matrix of pairwise distances estimated using the Maximum Composite Likelihood (MCL) approach, and then selecting the topology with superior log likelihood value. The tree is drawn to scale, with branch lengths measured in the number of substitutions per site. The analysis involved (*A*) complete mitochondrial (Mt) genome sequences of salamanders, toads and frogs (classified as *Rana* (blue highlight) or *Lithobates* (yellow highlight). Analysis of Mt genes (*B*) *cyb*, (*C*) *rnr1*, and (*D*) *rnr2* of selected frog species. Position of *R. catesbeiana* indicated by an asterisk. Evolutionary analyses were conducted in MEGA7 (Kumar et al. 2016).

### Comparative genomics

Estimates of the genomic sequence identity between of *N. parkeri*, *X. tropicalis* and *R.catesbeiana* were performed. We did this using Bloom filters (Bloom 1970), probabilistic data structures with bit sets for each genome's k-mers (collection of all subsequences of length k). In a previous study, this method was shown to provide concordant estimates of the genome sequence divergence of known model organisms (human and apes), and was applied to conifer genomes (Warren et al. 2015b).

This method is designed to compare the k-mer spectra of any two genomes by computing the k-mer set bit intersections of their respective Bloom filters. It is assumed that differences between the genomes are independently distributed. We point out that this method does not factor size differences in the genomes, nor structural rearrangements; instead it reports on the commonality over short sequence stretches. The reliability of the method also comes into question for very divergent genomes, as common k-mers are rarer, and precise values of sequence identity are not expected. As such, divergence figures are likely an underestimate of their true separation.

At k=25 bp, *R. catesbeiana* shares higher sequence identity (86.0 +/− 3.3×10^−4^ %) with the High Himalaya frog (*N. parkeri*) than to the more evolutionarily diverged *X. tropicalis* (79.2 +/− 3.4×10^−4^ %), which is estimated to have shared a common ancestor with bullfrog more than 200 MYA, more than twice as much (89 MYA) as between *N. parkeri* and *R. catesbeiana* (Table 3).

**Table 3.** ABySS-Bloom sequence identity calculations between various draft genomeassemblies^*^.

|  |  | Estimated time since divergence (MYA) |  |  |
| --- | --- | --- | --- | --- |
|  |  | *R. catesbeiana* | *N. parkeri* | *X. tropicalis* |
| Estimated identity (%) | *R. catesbeiana* |  | 89.0 | 208.6 |
| | *N. parkeri* | $86.01 \pm 3.32 \times 10^{-4}$ | | 208.6 |
| | *X. tropicalis* | $79.16 \pm 3.42 \times 10^{-3}$ | $77.92 \pm 2.51 \times 10^{-3}$ | |

\*k=25, Sequences $\geq 500$ bp

## Discussion

Amphibians are the only group where most of its members exhibit a life cycle that includes distinct independent aquatic larval and terrestrial juvenile/adult phases. The transition between the larval and juvenile phases requires substantial or complete remodeling of the organism (metamorphosis) in anticipation of a terrestrial lifestyle. Thus, this places amphibians in a unique position for the assessment of toxicological effects in both aquatic and terrestrial environments. As a model for human and mammalian perinatal development, including the transition from the aquatic environment of the womb to the outside world, *R. catesbeiana* is a preferable model to the *Xenopus* species, because of similar physiological transformations. In contrast, *Xenopus* remain aquatic throughout life (Helbing 2012).

Our sequence divergence analysis with the recently released *Xenopus* genome of a diploid species, *X. tropicalis* (version 9.0) highlights the reason why Pipids are generally unsuitable genome references for Ranid species due to their considerable evolutionary divergence. Our present work is consistent with earlier estimates of divergence dating over ~200 MYA. The initial assembly of another *Xenopus* species’ genome – that of the allotetraploid *X. laevis* inbred ‘J’ strain – has just been published (Session et al. 2016). The haploid genomes of these species are substantially smaller (1.7 and 3.1 Gbp, respectively) than the typical Ranid genome (Gregory 2017). Despite the difference in relative sizes, the number of high confidence predicted protein-encoding genes in *R. catesbeiana* (~22,000) is comparable (24,022 in *X.laevis*; ~20,000 in *X. tropicalis*) (Hellsten et al. 2010; Session et al. 2016). However, the sequence divergence at the nucleic acid level confirms the empirical challenges of using *Xenopus* genomes as scaffolds for RNA-Seq experiments in Ranids. Even the recently published *N. parkeri* genome (Sun et al. 2015), is substantially separated from *R. catesbeiana* by nearly 90 million years of evolutionary time (timetree.org).

The taxonomic classification of Ranid species is contentious and highly debated amongst scholars (Pauly et al., 2009). This stems largely from the suggested reclassification of the genus *Rana* in favor of *Lithobates* a few years ago (Frost et al. 2006). Our phylogenetic analyses based on comparisons at the nucleotide level of complete mitochondrial genomes and genes from selected salamanders, toads, and Ranids does not offer a rationale for the proposed change. It instead supports a close relationship between species classified as *Lithobates*, which may be considered a subgenus within the *Rana* genus. This observation is consistent with the recent phylogenetic analysis by Yuan et al. (2016).

The molecular mechanisms of amphibian metamorphosis have been predominantly studied using *X. laevis* and *X. tropicalis*, likely in no small part because of their amenability to captive breeding, which ensures a ready supply of research specimens (Parker et al. 1947). However, *Xenopus* larvae are typically much smaller than those of *R. catesbeiana*, with the consequence that each individual animal yields a smaller quantity of tissue for analysis. Indeed, *R.catesbeiana* tadpoles are large enough that techniques such as the cultured tail fin (C-fin) assay are possible, where multiple tissue biopsies are collected from an individual animal and cultured *ex vivo* in a variety of hormone or chemical conditions (Hinther et al. 2010; Hinther et al. 2011; Hinther et al. 2012; Hammond et al. 2013). With this assay design each animal is effectively exposed simultaneously and independently to every condition in the experiment, and the result of these different conditions can be evaluated within each individual animal using powerful repeated-measures statistics.

Analysis of TH-induced changes in tadpole back skin gene expression revealed an unprecedented view of the activation of new gene expression programs as an integral part of the transition from larval to adult skin. It is notable that the vast majority of annotated transcripts functioned in RNA/DNA processing roles. The wave of new transcripts that results from TH treatment must be spliced, capped with 7-methylguanosine, and polyadenylated. As the level of circulating TH increases in the tadpole, expression of key cell cycle control genes changes to regulate the proliferation of skin cells, including cyclin C and cyclin B (Buchholz et al. 2007; Skirrow et al. 2008; Suzuki et al. 2009). This is in contrast to TH-induced remodeling of the liver tissue, where these cellular processes were not as prominently represented (Birol et al. 2015).

Current lncRNA databases are mostly populated with sequences that were derived from human and mouse (Bu et al. 2012; Quek et al. 2015; Volders et al. 2015). Recent studies involving bovine (Huang et al. 2012; Weikard et al. 2013; Billerey et al. 2014), chicken (Li et al. 2012), pig (Zhao et al. 2015), diamondback moth (Etebari et al. 2015), and goat (Ren et al. 2016) have further contributed in enriching animal lncRNA datasets. A few databases exploring interactions of lncRNAs with proteins, RNAs, and viruses have also been developed, but mainly describing interactions in human (Zhang et al. 2014). In addition to expanding the effectiveness of RNA-Seq analyses through annotation of protein-encoding transcripts, the present study identified over 6,000 putative lncRNA candidates in the bullfrog. Despite being non-coding with relatively low level of sequence conservation, some lncRNAs contribute to structural organization, function, and evolution of genomes (Derrien et al. 2012). Examples include classical lncRNAs, such as X-inactive specific transcript (XIST), HOX transcript antisense RNA (HOTAIR), telomerase RNA component (TERC), and many more with roles in imprinting genomic loci (Thakur et al. 2004), transcription (Feng et al. 2006), splicing (Yan et al. 2005), translation (Wang et al. 2005), nuclear factor trafficking (Willingham et al. 2005), chromatin modification, shaping chromosome conformation, and allosterically regulating enzymatic activity (Ponting et al. 2009; Rinn and Chang 2012). These functional roles overlap with the major gene ontologies associated with the protein coding genes differentially expressed in the T3-treated back skin.

Dynamic regulation of lncRNAs has previously been observed during embryogenesis in *X. tropicalis* (Forouzmand et al. 2016). Additional studies have ascribed roles of lncRNAs in differentiation of mouse embryonic stem cells (Dinger et al. 2008), and shown tissue specific patterns of expression in human tissues (Cabili et al. 2011). HOTAIR expression is transcriptionally induced by estradiol in the MCF7 breast cancer cell line, and its promoter contains multiple estrogen response elements (Bhan et al. 2013). As the candidate lncRNAs that we identified represented 1/6^th^ of the differentially expressed genes in response to TH treatment, this suggests an important role for lncRNA in the amphibian metamorphic gene expression program initiated by this hormone. The data presented herein extend hormonal regulation of lncRNAs to postembryonic developmental processes. Further experiments to elucidate their role in this context are ongoing.

The bullfrog genome is larger than those of *N. parkeri, X. laevis*, and *X. tropicalis*, in concordance with earlier predictions for many Ranid genomes (Mazin 1980; Buisine et al. 2015; Sun et al. 2015; Session et al. 2016). Unlike *X. laevis*, this does not appear to be due to an allotetraploidization event in the Ranid progenitor species (Wang et al. 2000; Zhu and Wang 2006). Another possibility for genome enlargement is integration of foreign DNA, which can manifest as DNA sequence repeat elements. Many of these integrations are likely to be derived from ancestral integration of viral genomes into the host genome (Belyi et al. 2010). The genomes of group I linear viruses, such as herpesviruses and FV3, include integrases that enable integration of their genetic material into that of the host cell during replication (Rohozinski and Goorha 1992). However, the limited alignment of a single putative FV3 US22 domain found in many animal DNA viruses and some vertebrates (Zhang et al. 2011) does not support the inclusion of a ranavirus genome into our assembled bullfrog genome *per se*, but may point to previous irreversible integration of viral genetic material in the evolutionary history of the bullfrog.

The *R. catesbeiana* genome presented herein provides an unprecedented resource for Ranidae. For example, it will inform the design and/or interpretation of high throughput transcriptome sequencing (RNA-Seq), chromatin immunoprecipitation sequencing (ChIP-Seq), and proteomics experiments. We anticipate that this resource will be valuable for conservation efforts such as identifying host/pathogen interactions and to identify environmental impacts of climate change and pollution on the development and reproduction of Ranid species worldwide.

## Methods

### Sample collection

Liver tissue was collected from an adult male *R. catesbeiana* specimen that had been caught in Victoria, BC, Canada and housed at the University of Victoria Outdoor Aquatics Unit. The tissue was taken under the appropriate sanctioned protocols and permits approved by the University of Victoria Animal Care Committee (Protocol #2015-028) and British Columbia Ministry of Forests, Lands and Natural Resource Operations (MFLNRO) permit VI11-71459. This frog was euthanized using 1% w/v tricaine methane sulfonate in dechlorinated municipal water containing 25 mM sodium bicarbonate prior to tissue collection. Dissected liver pieces were preserved in RNA*later* (Thermo Fisher Scientific Inc., Waltham, MA, USA) at room temperature followed by incubation at 4°C for 24 h. Tissue samples were subsequently moved to storage at −20°C prior to DNA isolation. Total DNA was isolated using the DNeasy Blood and Tissue Kit (QIAGEN Inc., Mississauga, ON, Canada; Cat# 69506) with the inclusion of RNase treatment as per the manufacturer’s protocol, and stored at −20 °C prior to library preparation.

### DNA library preparation and sequencing

All reagent kits used were from the same vendor (Illumina, San Diego, CA) unless otherwise stated. Two sets of paired-end tag (PET) libraries were constructed: 1) sixteen libraries were produced using 1 μg of DNA using custom NEBNext DNA Library Prep Reagents from New England BioLabs Canada (Whitby, ON); and 2) four libraries were constructed using 0.5 μg of DNA and the custom NEB Paired-End Sample Prep Premix Kit (New England BioLabs Canada). DNA sequence reads were generated from these libraries according to the manufacturer’s instructions on the Illumina HiSeq 2000 platform (Illumina, San Diego, CA) in “High Throughput” mode with the HiSeq SBS Kit v3, on the Illumina HiSeq 2500 platform in “Rapid” mode with the HiSeq Rapid SBS kit v1, or on the Illumina MiSeq platform with the MiSeq Reagent Kit v2. See Table 2 for additional details.

**Table 2.** Sequencing data for *R. catesbeiana* genome assembly

| Library protocol | Read length (bp) | Sequencing platform | Nominal fragment length (bp) | # Libraries | # Reads (M) | Fold coverage |
| --- | --- | --- | --- | --- | --- | --- |
| PET | 150 | HiSeq 2000 | 600 | 8 | 1,187 | 30 |
| PET | 250 | HiSeq 2500 | 550 | 4 | 736 | 31 |
| PET | 500 | MiSeq | 600 | 8 (same) | 66 | 5 |
| MPET | 100 | HiSeq 2000 | 9,000- 13,000 | 4 | 262 | N/A |
| SLR | N/A | HiSeq 2000 | 0.5 - 14,000 | 1 | 0.6 | 0.5 |

The mate pair (MPET, a.k.a. jumping) libraries were constructed using 4 μg of DNA and the Nextera Mate Pair Library Preparation Kit, according to the manufacturer’s protocol. One-hundred bp paired-end reads were generated on the Illumina HiSeq 2000 platform with the HiSeq SBS Kit v3. The Synthetic Long-Read (SLR, a.k.a. Moleculo) library was constructed using 500 ng DNA and Illumina’s TruSeq SLR library prep kit with 8 – 10 kb size DNA fragments. Libraries were loaded on an Illumina HiSeq 2500 platform for 125 bp paired end sequencing.

Combined, this approach accounted for 66-fold sequence coverage of the approximately 6 Gbp bullfrog genome (Table 2).

### Computing hardware

Sequence assemblies were performed on high performance computer clusters located at the Canada’s Michael Smith Genome Sciences Centre, and consisted of nodes with 48 GB of RAM and dual Intel Xeon X-5650 2.66GHz CPUs running Linux CentOS 5.4 or 128 GB RAM and dual Intel Xeon E5-2650 2.6 GHz CPUs running Linux CentOS 6. Computational analyses used either this hardware, or nodes consisting of 24 GB of RAM and dual Intel Xeon X-5550 2.67 GHz CPUs running Red Hat Enterprise Linux 5 as part of WestGrid, Compute Canada.

### Read merging

PET read pairs were merged sequentially using the ABySS-mergepairs tool (Birol et al. 2013) and Konnector (version 1.9.0) (Vandervalk et al. 2015). Bloom filters were constructed from all reads using the ABySS-Bloom utility (Warren et al. 2015b), and every tenth value of k between 75 and 245 bp, inclusive. Reads from potentially mixed clusters on the sequencing flow cells (determined by the Illumina chastity flag) were discarded, and the remaining reads were trimmed to the first base above a quality threshold (Q=3 on the phred scale) prior to merging.

### Assembly process

ABySS (version 1.9.0) was used to reconstruct the *R. catesbeiana* genome (Simpson et al. 2009). For the initial sequence assembly, three sets of reads were used: (i) merged reads described above from paired-end Illumina HiSeq 150 bp, 250 bp, and MiSeq 500 bp libraries, (ii) unmerged reads from these same libraries, and (iii) synthetic long-reads. The unmerged HiSeq and MiSeq PET reads were also used for paired linking information to generate contigs. Finally, the MPET reads were used to bridge over regions of repetitive sequence to form scaffolds (see Table 2 for summary statistics of the sequencing data).

A certain fraction of the unresolved bases within these scaffolds were recovered using Sealer version 1.9.0 (Paulino et al. 2015). Sealer uses a Bloom filter representation of a de Bruijn graph constructed using k-mers derived from the genomic reads to find a path between the sequences flanking scaffold gaps, and fill in the consensus sequence. In comparison to the fixed k-mer length of the whole genome assembly method, it uses a range of k-mer lengths to navigate repeat and low coverage areas within the graph. The Bloom filters that were used during the read merging phase were reused, and default values were used for all parameters except for “--flank-length=260” and “--max-paths=10”.

The resulting ABySS scaffold assembly (k = 160 bp) was rescaffolded with RAILS version 0.1 (Warren 2016) (ftp://ftp.bcgsc.ca/supplementary/RAILS-d250-i0.99) using both SLR data and Kollector (Kucuk, et al., *in press*) targeted gene reconstructions (TGA; Supplemental Table S1). In RAILS, long sequences are aligned against a draft assembly (BWA-MEM V0.7.13-r1126 (Li 2013)-a-t16), and the alignments are parsed and inspected, tracking the position and orientation of each in assembly draft sequences, satisfying minimum alignment requirements (at minimum 250 anchoring bases with 99% sequence identity or more used in this study). Sequence scaffolding is performed using the scaffolder algorithms from LINKS (Warren et al. 2015a), modified to automatically fill gaps with the sequence that informed the merge. The resulting assembly was sequentially rescaffolded with a composite reference transcriptome (Bullfrog Annotation Resource for the Transcriptome; BART, Supplemental Table S1; see below) with ABySS-longseqdist v1.9.0 (l=50, S=1000-) (Warren et al. 2015b). It was further rescaffolded iteratively with LINKS (v1.7) using a variety of long sequence data (Supplemental Table S1) including SLR data (10 iterations-d 1-10:1 kbp,-t 10-1:-1,-k 20), MPET (-k 20,-t 5,- d 7.1 kbp,-e 0.9) and other assembly draft data (Kollector targeted reconstructions and whole genome assembly at k=128 bp combined, 7 iterations-d 1-15:2.5 kbp,-t 20,10,5,5,4,4,4 k=20).

A final round of automated gap closing in the assembled draft was performed by Sealer. Completeness of the assembly was evaluated by comparison to a set of ultra-conserved core eukaryotic genes (Parra et al. 2009).

### Protein coding gene prediction

The MAKER2 genome annotation pipeline (version 2.31.8) was used to predict genes in the draft *R. catesbeiana* genome (Holt and Yandell 2011). This framework included RepeatMasker (Smit et al. 2013) to mask repetitive sequence elements based on the core RepBase repeat library (Jurka et al. 2005). Augustus (Stanke et al. 2006), SNAP (Korf 2004) and GeneMark (Ter-Hovhannisyan et al. 2008) were also run within the MAKER2 pipeline to produce *ab initio* gene predictions. BLASTx (Camacho et al. 2009), BLASTn (Camacho et al. 2009), and exonerate (Slater and Birney 2005) alignments of human and amphibian Swiss-Prot protein sequences (retrieved 16 February 2016) (UniProt 2015) and BART were combined with the gene predictions to yield the gene models. MAKER2 was first applied to an early version of the bullfrog genome assembly, and the resulting gene models used for retraining as described in (Cantarel et al. 2008).

We refined MAKER2’s predicted gene list further by identifying a high confidence set, better suited for downstream biological analyses. Three criteria were considered: 1) the predicted transcripts must have at least one splice site, and all putative splice sites must be confirmed by an alignment to external transcript evidence; 2) the coding DNA sequence (CDS) of each transcript must have a BLASTn alignment to a BART contig with at least 95% identity along 99% of its length; or 3) the protein sequence encoded by the CDS must have a BLASTp (Camacho et al. 2009) alignment to a human or amphibian Swiss-Prot protein sequence with at least 50% identity along 90% of its length (Supplemental Fig. S1).

### Functional annotation

The high confidence set of transcripts was annotated according to the best BLASTp alignment of each putative encoded protein to the Swiss-Prot database (UniProt 2015), when available. There were two levels of confidence for the annotations, 1) the most robust were identified using GeneValidator, which compares protein coding gene predictions with similar database proteins (Dragan et al. 2016), where those having a score of 90 or greater were definitively identified as the Swiss-Prot sequence they aligned to; and 2) all other annotated transcripts were considered to encode “hypothetical” proteins similar to their best Swiss-Prot hit, provided that they aligned with at least 25% identity along 50% of their length.

### Construction of a composite reference transcriptome (Bullfrog Annotation Resource for the Transcriptome; BART)

Transcriptome assemblies were generated from 32 *R. catesbeiana* tadpole samples (representing 5 tissues under several different chemical and temperature exposure conditions) using Trans-ABySS (Robertson et al. 2010) (see Supplemental Table S5). The transcripts from each independent assembly were aligned using the BLAST-like Alignment Tool (Kent 2002) or parallelized BLAT (pblat; icebert.github.io/pblat/) to identify highly similar sequences, where only the longest example of each set of similar sequences was retained. This process produced 794,291 transcripts 500 bp or longer, collected and termed the Bullfrog Annotation Resource for the Transcriptome (BART). Further, we report 1,341,707 transcripts between 200 and 499 bp long (termed BART Jr.).

### IncRNA prediction

To complement the protein coding gene predictions, a computational pipeline was developed to identify putative lncRNAs in the *R. catesbeiana* composite reference transcriptome BART. As there is a paucity of conserved sequence features that may positively identify lncRNA transcripts, we instead took a subtractive approach, and omitted transcripts that were predicted to have coding potential or had sequence similarity to known protein encoding transcripts, as has been advocated in previous studies (Tan et al. 2013; Etebari et al. 2015). See Supplemental Methods for additional details.

We then used CD-HIT-EST (Fu et al. 2012) (v4.6.6,-c 0.99) to identify and remove contigs with significantly redundant sequence content. The remaining transcripts were then interrogated for the presence of a poly(A) tail and one of 16 polyadenylation signal hexamer motifs (see Supplemental Table S6). The contigs were aligned to the genome assembly using GMAP (v2016-05-01,-f samse,-t 20) (Wu and Watanabe 2005), and instances where there was a 3’ sequence mismatch due to a run of As, or a 5’ mismatch due to a run of Ts (in cases where the strand specific sequencing failed, and a RNA molecule complementary to the actual transcript was sequenced) prompted a search for the presence of a hexamer motif within 50 bp upstream (relative to the direction of coding) of the putative transcript cleavage site. Contigs containing a poly(A) tail and a hexamer motif were selected for further analysis. We are aware that not all lncRNA are polyadenylated. The poly(A) tail filter was put in place to reduce the proportion of spurious transcripts, retained introns and assembly artifacts.

Candidate lncRNA transcripts were aligned to the draft genome with GMAP (version 2015-12-31,-f 2,-n 2, --suboptimal-score=0, --min-trimmed-coverage=0.9, --min-identity=0.9) (Wu and Watanabe 2005), and those that had at least 90% of their sequence identified across no more than two separate genomic scaffolds with 90% sequence identity were retained. Alignments where the exon arrangement was not collinear with the original contig sequence were omitted. Further evidence of conservation of lncRNA candidates among amphibian species was obtained using a comprehensive amphibian transcriptome shotgun assembly database, as described in the Supplemental Methods and Supplemental Table S7.

### Differential gene expression analysis

As an example of the utility of the draft genome assembly and high confidence gene predictions, RNA-Seq reads from six premetamorphic *R. catesbeiana* tadpoles exposed to 10 pmol/g body weight T3 or dilute sodium hydroxide vehicle control for 48 h were used to characterize the T3-induced gene expression program in the back skin (Supplemental Methods; Supplemental Table S5). The 100 bp paired-end reads were aligned to the draft genome using STAR (version 2.4.1d, --alignIntronMin 30, --alignIntronMax 500000, --outFilterIntronMotifs RemoveNoncanonicalUnannotated, --outFilterMismatchNMax 10, --sjdbOverhang 100) (Dobin et al. 2013), and read counts per transcript were quantified using HTSeq (version 0.6.1, default settings) (Anders et al. 2015). Differential expression in response to T3 treatment was assessed using the DESeq2 software package (version 1.10.1, alpha=0.05) (Love et al. 2014), and significance was considered where the Benjamini – Hochberg adjusted p-value was less than 0.05. Transcripts with zero counts in all six samples were excluded from the analysis. qPCR analysis of transcripts is described in the Supplemental Methods.

### Gene ontology (GO) and pathway analysis

Due to the particularly extensive biological information available for human proteins, a second round of BLASTp alignments were performed between the high confidence set of predicted proteins and the Swiss-Prot human proteins, using the same alignment thresholds noted above. The Uniprot accession IDs and log fold-changes of the differentially expressed genes were collected, input to the IngenuityPathway Analysis tool (Qiagen Bioinformatics, Redwood City, CA), and its core analysis was run with default settings. The Database for Annotation, Visualization and Integrated Discovery (DAVID) v6.8 (Huang et al. 2009) was also used with default settings to perform gene annotation enrichment analysis on the differentially expressed genes versus the background of all bullfrog genes with Uniprot annotations. The enriched annotations were visualized with ReviGO (Supek et al. 2011) with default settings.

### Iridovirus integration experiment

The FV3 ranavirus genome and associated protein coding DNA sequences were downloaded from NCBI (GenBank Accession NC_005946.1), and aligned to the assembly with GMAP version 2015-12-31 (Wu and Watanabe 2005) using default settings.

### Mitochondrial genome assembly and finishing

The Mitochondrial (Mt) genome sequence was identified integrally in our whole genome assembly. Rounds of scaffolding effectively brought unincorporated Mt contigs to the edge of the scaffold, and after inspection were removed by breaking the redundant scaffolds at N’s. Multiple sequence alignments were done between our sequence, two NCBI references originating from China (GenBank accessions NC02296 and KF049927.1) and one from Japan (AB761267), using MUSCLE (Edgar 2004) from the MEGA phylogenetic analysis package (Kumar et al. 2016) using default values. These analyses indicated that the Mt sequence reported herein is most similar to the Japanese sequence, but with some discrepancies in two repeat regions, namely the absence of a 161 bp sequence at coordinate 15,270, and a 12-bp insertion at coordinate 9,214 relative to AB761267. We resolved these misassemblies using the correct Japanese reference sequence for these regions as candidates for a targeted *de novo* assembly of Illumina paired-end 250 bp reads with TASR (Warren and Holt 2011); v1.7 with-w 1-i 1-k 15-m 20-o 2-r 0.7. TASR uses whole reads for assemblies, and mitigates misassemblies otherwise caused by breaking reads into k-mers. The resulting TASR contigs that captured the correct sequences were inserted into our assembly in the corresponding regions. Transfer RNA (tRNA) and protein coding genes were annotated by GMAP alignment of the gene sequences included in KF049927.1.

### Phylogenetic analyses

Complete mitochondrial genome sequences of selected salamanders and frogs (Supplemental Table S8) were compared using the MEGA phylogenetic package (Kumar et al. 2016). In another set of phylogenies, we also compared the mitochondrial genes *cyb* and 12S and 16S rRNA *rnr1* and *rnr2* of selected amphibian species (Supplemental Table S9). For these analyses, we first generated multiple sequence alignments of the genome or gene sequences described above using either MUSCLE (Edgar 2004) or clustalw (Chenna et al. 2003) (v1.83 with gap opening and extension penalty of 15 and 6.66 for both the pairwise and multiple alignment stages, DNA weight matrix IUB with transition weight of 0.5), and used the resulting pairwise alignments as input for MEGA7 (Kumar et al. 2016). The evolutionary history was inferred by using the Maximum Likelihood method based on the Tamura-Nei model (Tamura and Nei 1993), where initial trees for the heuristic search are obtained by applying Neighbor-Join and BioNJ algorithms to a matrix of pairwise distances estimated using the Maximum Composite Likelihood approach, and then selecting the topology with superior log likelihood value.

### Comparative genome analysis using Bloom filters

The genomes of *N. parkeri* (version 2; http://dx.doi.org/10.5524/100132), *X. tropicalis* (version 9.0, downloaded from xenbase.org) and *R. catesbeiana* (the present study) were compared for their k-mer (k=25) contents using ABySS-Bloom, a utility designed to approximate sequence divergence between draft genomes (Warren et al. 2015b).

### Repetitive sequence element detection

The content of repetitive sequence elements in the draft genome assembly was evaluated with RepeatMasker (version 4.0.6) (Smit et al. 2013) with default settings. The RepBase collection of repeat sequence elements was supplemented with novel elements identified using RepeatModeler (version 1.0.8) (Smit and Hubley 2008) with RMBlast (version 2.2.27+, http://www.repeatmasker.org/RMBlast.html) applied to the draft genome assembly with default settings.

### Data Access

The whole genome sequence data and final assembly of the North American bullfrog genome with annotated MAKER2 gene predictions is available at NCBI-Genbank under accession (LIAG00000000, BioProject PRJNA285814).

The Mt genome assembly was submitted to NCBI GenBank under accession KX686108. The RNA-Seq reads and assembled BART contigs are available under NCBI BioProject PRJNA286013, and the BART Jr. contigs (collection of short transcripts between 200 and 499 bp in length in the composite transcriptome that forms the basis for BART) are available from the BCGSC ftp:ftp://ftp.bcgsc.ca/supplementary/bullfrog. The candidate lncRNA sequences and genomic coordinates are also available from the BCGSC ftp.

## Acknowledgements

This work has been supported by Genome British Columbia and Natural Sciences and Engineering Research Council of Canada, with additional support provided by the National Human Genome Research Institute of the National Institutes of Health (under award number R01HG007182). The content is solely the responsibility of the authors, and does not necessarily represent the official views of the National Institutes of Health or other funding organizations. We thank Dr. Belaid Moa for advanced research computing support from WestGrid, Compute Canada and the University of Victoria computing systems.

## References

AmphibiaWeb. 2017. Worldwide Amphibian Declines: How big is the problem, what are the causes and what can be done? Available at: http://www.amphibiaweb.org. [accessed January 4].

Anders S, Pyl PT, Huber W. 2015. HTSeq-a Python framework to work with high-throughput sequencing data. Bioinformatics 31(2):166–169.

Beaudoing E, Freier S, Wyatt JR, Claverie JM, Gautheret D. 2000. Patterns of variant polyadenylation signal usage in human genes. Genome res 10(7):1001–1010.

Belyi VA, Levine AJ, Skalka AM. 2010. Unexpected inheritance: multiple integrations of ancient bornavirus and ebolavirus/marburgvirus sequences in vertebrate genomes. PLoS pathogens 6(7):e1001030.

Bhan A, Hussain I, Ansari KI, Kasiri S, Bashyal A, Mandal SS. 2013. Antisense transcript long noncoding RNA (lncRNA) HOTAIR is transcriptionally induced by estradiol. J mol biol 425(19):3707–3722.

Billerey C, Boussaha M, Esquerré D, Rebours E, Djari A, Meersseman C, Klopp C, Gautheret D, Rocha D. 2014. Identification of large intergenic non-coding RNAs in bovine muscle using next-generation transcriptomic sequencing. BMC genomics 15:499.

Birol I, Behsaz B, Hammond SA, Kucuk E, Veldhoen N, Helbing CC. 2015. De novo transcriptome assemblies of Rana (Lithobates) catesbeiana and Xenopus laevis tadpole livers for comparative genomics without reference genomes. PloS one 10(6):e0130720.

Birol I, Raymond A, Jackman SD, Pleasance S, Coope R, Taylor GA, Yuen MM, Keeling CI, Brand D, Vandervalk BP et al. 2013. Assembling the 20 Gb white spruce (*Picea glauca*) genome from whole-genome shotgun sequencing data. Bioinformatics 29(12): 1492–1497.

Bloom BH. 1970. Space/time trade-offs in hash coding with allowable errors. Commun Acm 13(7):422–426.

Bu D, Yu K, Sun S, Xie C, Skogerbo G, Miao R, Xiao H, Liao Q, Luo H, Zhao G et al. 2012. NONCODE v3.0: integrative annotation of long noncoding RNAs. Nucleic acids res 40:D210–215.

Buchholz DR, Heimeier RA, Das B, Washington T, Shi YB. 2007. Pairing morphology with gene expression in thyroid hormone-induced intestinal remodeling and identification of a core set of TH-induced genes across tadpole tissues. Dev biol 303(2):576–590.

Buisine N, Ruan X, Bilesimo P, Grimaldi A, Alfama G, Ariyaratne P, Mulawadi F, Chen J, Sung WK, Liu ET et al. 2015. *Xenopus tropicalis* genome re-scaffolding and re-annotation reach the resolution required for *in vivo* ChIA-PET analysis. PloS one 10(9):e0137526.

Cabili MN, Trapnell C, Goff L, Koziol M, Tazon-Vega B, Regev A, Rinn JL. 2011. Integrative annotation of human large intergenic noncoding RNAs reveals global properties and specific subclasses. Genes dev 25(18):1915–1927.

Camacho C, Coulouris G, Avagyan V, Ma N, Papadopoulos J, Bealer K, Madden TL. 2009. BLAST+: architecture and applications. BMC bioinformatics 10:421.

Cantarel BL, Korf I, Robb SM, Parra G, Ross E, Moore B, Holt C, Sanchez Alvarado A, Yandell M. 2008. MAKER: an easy-to-use annotation pipeline designed for emerging model organism genomes. Genome res 18(1):188–196.

Chenna R, Sugawara H, Koike T, Lopez R, Gibson TJ, Higgins DG, Thompson JD. 2003. Multiple sequence alignment with the Clustal series of programs. Nucleic acids res 31(13):3497–3500.

Clark K, Karsch-Mizrachi I, Lipman DJ, Ostell J, Sayers EW. 2016. GenBank. Nucleic acids res 44(D1):D67–72.

Derrien T, Johnson R, Bussotti G, Tanzer A, Djebali S, Tilgner H, Guernec G, Martin D, Merkel A, Knowles DG et al. 2012. The GENCODE v7 catalog of human long noncoding RNAs: analysis of their gene structure, evolution, and expression. Genome res 22(9):1775–1789.

Dinger ME, Amaral PP, Mercer TR, Pang KC, Bruce SJ, Gardiner BB, Askarian-Amiri ME, Ru K, Soldà G, Simons C et al. 2008. Long noncoding RNAs in mouse embryonic stem cell pluripotency and differentiation. Genome res 18(9):1433–1445.

Dobin A, Davis CA, Schlesinger F, Drenkow J, Zaleski C, Jha S, Batut P, Chaisson M, Gingeras TR. 2013. STAR: ultrafast universal RNA-seq aligner. Bioinformatics 29(1):15–21.

Dragan MA, Moghul I, Priyam A, Bustos C, Wurm Y. 2016. GeneValidator: identify problems with protein-coding gene predictions. Bioinformatics 32(10):1559–1561.

Edgar RC. 2004. MUSCLE: multiple sequence alignment with high accuracy and high throughput. Nucleic acids res 32(5):1792–1797.

Eggert C. 2004. Sex determination: the amphibian models. Reprod nutr dev 44(6):539–549.

Etebari K, Furlong MJ, Asgari S. 2015. Genome wide discovery of long intergenic non-coding RNAs in Diamondback moth (*Plutella xylostella*) and their expression in insecticide resistant strains. Sci rep 5:14642.

Feng J, Bi C, Clark BS, Mady R, Shah P, Kohtz JD. 2006. The Evf-2 noncoding RNA is transcribed from the Dlx-5/6 ultraconserved region and functions as a Dlx-2 transcriptional coactivator. Genes dev 20(11):1470–1484.

Finn RD, Bateman A, Clements J, Coggill P, Eberhardt RY, Eddy SR, Heger A, Hetherington K, Holm L, Mistry J et al. 2014. Pfam: the protein families database. Nucleic acids res 42:D222–230.

Finn RD, Clements J, Arndt W, Miller BL, Wheeler TJ, Schreiber F, Bateman A, Eddy SR. 2015. HMMER web server: 2015 update. Nucleic acids res 43(W1):W30–38.

Forouzmand E, Owens ND, Blitz IL, Paraiso KD, Khokha MK, Gilchrist MJ, Xie X, Cho KW. 2016. Developmentally regulated long non-coding RNAs in *Xenopus tropicalis*. Dev biol. (in press). doi:10.1016/j.ydbio.2016.06.016.

Frost DR, Grant T, Faivovich J, Bain RH, Haas A, Haddad CFB, De Sa RO, Channing A, Wilkinson M, Donnellan SC et al. 2006. The amphibian tree of life. Bull Am Mus Nat Hist 297:1–370.

Fu L, Niu B, Zhu Z, Wu S, Li W. 2012. CD-HIT: accelerated for clustering the next-generation sequencing data. Bioinformatics 28(23):3150–3152.

Gregory TR. 2017. Animal Genome Size Database. Available at: http://www.genomesize.com. [accessed January 4].

Hammond SA, Carew AC, Helbing CC. 2013. Evaluation of the effects of titanium dioxide nanoparticles on cultured *Rana catesbeiana* tailfin tissue. Front genet 4:251.

Hayes TB, Falso P, Gallipeau S, Stice M. 2010. The cause of global amphibian declines: a developmental endocrinologist's perspective. J exp biol 213(6):921–933.

Helbing CC. 2012. The metamorphosis of amphibian toxicogenomics. Front genet 3:37.

Hellsten U, Harland RM, Gilchrist MJ, Hendrix D, Jurka J, Kapitonov V, Ovcharenko I, Putnam NH, Shu S, Taher L et al. 2010. The genome of the Western clawed frog *Xenopus tropicalis*. Science 328(5978):633–636.

Hinther A, Bromba CM, Wulff JE, Helbing CC. 2011. Effects of triclocarban, triclosan, and methyl triclosan on thyroid hormone action and stress in frog and mammalian culture systems. Environ sci technol 45(12):5395–5402.

Hinther A, Domanski D, Vawda S, Helbing CC. 2010. C-fin: a cultured frog tadpole tail fin biopsy approach for detection of thyroid hormone-disrupting chemicals. Environ toxicol chem 29(2):380–388.

Hinther A, Edwards TM, Guillette LJ, Jr., Helbing CC. 2012. Influence of nitrate and nitrite on thyroid hormone responsive and stress-associated gene expression in cultured *Rana catesbeiana* tadpole tail fin tissue. Front genet 3:51.

Holt C, Yandell M. 2011. MAKER2: an annotation pipeline and genome-database management tool for second-generation genome projects. BMC bioinformatics 12:491.

Huang DW, Sherman BT, Lempicki RA. 2009. Systematic and integrative analysis of large gene lists using DAVID bioinformatics resources. Nat protoc 4(1):44–57.

Huang W, Long N, Khatib H. 2012. Genome-wide identification and initial characterization of bovine long non-coding RNAs from EST data. Anim genet 43(6):674–682.

Jurka J, Kapitonov VV, Pavlicek A, Klonowski P, Kohany O, Walichiewicz J. 2005. Repbase Update, a database of eukaryotic repetitive elements. Cytogenet genome res 110(1–4):462–467.

Kamachi Y, Kondoh H. 2013. Sox proteins: regulators of cell fate specification and differentiation. Development 140(20):4129–4144.

Kent WJ. 2002. BLAT--the BLAST-like alignment tool. Genome res 12(4):656–664.

Kiemnec-Tyburczy KM, Richmond JQ, Savage AE, Lips KR, Zamudio KR. 2012. Genetic diversity of MHC class I loci in six non-model frogs is shaped by positive selection and gene duplication. Heredity 109(3):146–155.

Kong L, Zhang Y, Ye ZQ, Liu XQ, Zhao SQ, Wei L, Gao G. 2007. CPC: assess the protein-coding potential of transcripts using sequence features and support vector machine. Nucleic acids res 35:W345–349.

Korf I. 2004. Gene finding in novel genomes. BMC bioinformatics 5:59.

Kucuk E, Chu J, Vandervalk BP, Hammond SA, Warren RL, Birol I. in press. Kollector: transcript-informed, targeted *de novo* assembly of gene loci. Bioinformatics.

Kumar S, Stecher G, Tamura K. 2016. MEGA7: molecular evolutionary genetics analysis version 7.0 for bigger datasets. Molecular biol evol 33(7):1870–1874.

Lesbarrères D, Balseiro A, Brunner J, Chinchar VG, Duffus A, Kerby J, Miller DL, Robert J, Schock DM, Waltzek T et al. 2012. Ranavirus: past, present and future. Biol Letters 8(4):481–483.

Li H. 2013. Aligning sequence reads, clone sequences and assembly contigs with BWA-MEM. arXiv:1303.3997v1301 [q-bio.GN]

Li T, Wang S, Wu R, Zhou X, Zhu D, Zhang Y. 2012. Identification of long non-protein coding RNAs in chicken skeletal muscle using next generation sequencing. Genomics 99(5):292–298.

Liu X, Li Y. 2009. Aquaculture enclosures relate to the establishment of feral populations of introduced species. PloS one 4(7):e6199.

Love MI, Huber W, Anders S. 2014. Moderated estimation of fold change and dispersion for RNA-seq data with DESeq2. Genome biol 15(12):550.

Maher SK, Wojnarowicz P, Ichu TA, Veldhoen N, Lu L, Lesperance M, Propper CR, Helbing AC. 2016. Rethinking the biological relationships of the thyroid hormones, L-thyroxine and 3,5,3’-triiodothyronine. Comp biochem physiol D 18:44–53.

Mallo M, Alonso CR. 2013. The regulation of Hox gene expression during animal development. Development 140(19):3951–3963.

Mazin AL. 1980. Amounts of nuclear-DNA in anurans of the USSR. Experientia 36(2):190–191.

Parker FJ, Robbins SL, Loveridge A. 1947. Breeding, rearing and care of the South African clawed frog (*Xenopus laevis*). Am Nat 81(796):38–49.

Parra G, Bradnam K, Ning Z, Keane T, Korf I. 2009. Assessing the gene space in draft genomes. Nucleic acids res 37(1):289–297.

Paulino D, Warren RL, Vandervalk BP, Raymond A, Jackman SD, Birol I. 2015. Sealer: a scalable gap-closing application for finishing draft genomes. BMC bioinformatics 16:230.

Pauly GB, Hillis DM, Cannatella DC. 2009. Taxonomic freedom and the role of official lists of species names. Herpetologica 65(2):115–128.

Ponting CP, Oliver PL, Reik W. 2009. Evolution and functions of long noncoding RNAs. Cell 136(4):629–641.

Quek XC, Thomson DW, Maag JL, Bartonicek N, Signal B, Clark MB, Gloss BS, Dinger ME. 2015. lncRNAdb v2.0: expanding the reference database for functional long noncoding RNAs. Nucleic acids res 43:D168–173.

Quinn JJ, Chang HY. 2016. Unique features of long non-coding RNA biogenesis and function. Nat rev genet 17(1):47–62.

Raffel TR, Romansic JM, Halstead NT, McMahon TA, Venesky MD, Rohr JR. 2013. Disease and thermal acclimation in a more variable and unpredictable climate. Nat Clim Change 3(2):146–151.

Ren H, Wang G, Chen L, Jiang J, Liu L, Li N, Zhao J, Sun X, Zhou P. 2016. Genome-wide analysis of long non-coding RNAs at early stage of skin pigmentation in goats (*Capra hircus*). BMC genomics 17:67.

Rinn JL, Chang HY. 2012. Genome regulation by long noncoding RNAs. Ann rev biochem 81:145–166.

Robertson G, Schein J, Chiu R, Corbett R, Field M, Jackman SD, Mungall K, Lee S, Okada HM, Qian JQ et al. 2010. *De novo* assembly and analysis of RNA-seq data. Nat methods 7(11):909–912.

Rohozinski J, Goorha R. 1992. A frog virus 3 gene codes for a protein containing the motif characteristic of the INT family of integrases. Virology 186(2):693–700.

Schmidt J, Piekarski N, Olsson L. 2013. Cranial muscles in amphibians: development, novelties and the role of cranial neural crest cells. J anat 222(1):134–146.

Session AM, Uno Y, Kwon T, Chapman JA, Toyoda A, Takahashi S, Fukui A, Hikosaka A, Suzuki A, Kondo M et al. 2016. Genome evolution in the allotetraploid frog *Xenopus laevis*. Nature 538(7625):336–343.

Simpson JT, Wong K, Jackman SD, Schein JE, Jones SJ, Birol I. 2009. ABySS: a parallel assembler for short read sequence data. Genome res 19(6):1117–1123.

Skirrow RC, Veldhoen N, Domanski D, Helbing CC. 2008. Roscovitine inhibits thyroid hormone-induced tail regression of the frog tadpole and reveals a role for cyclin C/Cdk8 in the establishment of the metamorphic gene expression program. Dev dyn 237(12):3787–3797.

Slater GS, Birney E. 2005. Automated generation of heuristics for biological sequence comparison. BMC bioinformatics 6:31.

Smit AFA, Hubley R. 2008. RepeatModeler Open-1.0

Smit AFA, Hubley R, Green P. 2013. RepeatMasker Open-4.0

Stanke M, Schoffmann O, Morgenstern B, Waack S. 2006. Gene prediction in eukaryotes with a generalized hidden Markov model that uses hints from external sources. BMC bioinformatics 7:62.

Sumida M, Kato Y, Kurabayashi A. 2004. Sequencing and analysis of the internal transcribed spacers (ITSs) and coding regions in the EcoR I fragment of the ribosomal DNA of the Japanese pond frog *Rana nigromaculata*. Genes genet sys 79(2):105–118.

Sun YB, Xiong ZJ, Xiang XY, Liu SP, Zhou WW, Tu XL, Zhong L, Wang L, Wu DD, Zhang BL et al. 2015. Whole-genome sequence of the Tibetan frog *Nanorana parkeri* and the comparative evolution of tetrapod genomes. Proc natl acad sci U S A 112(11): E1257–1262.

Supek F, Bošnjak M, Škunca N, Šmuc T. 2011. REVIGO summarizes and visualizes long lists of gene ontology terms. PloS one 6(7):e21800.

Suzek BE, Wang Y, Huang H, McGarvey PB, Wu CH, UniProt C. 2015. UniRef clusters: a comprehensive and scalable alternative for improving sequence similarity searches. Bioinformatics 31(6):926–932.

Suzuki K, Machiyama F, Nishino S, Watanabe Y, Kashiwagi K, Kashiwagi A, Yoshizato K. 2009. Molecular features of thyroid hormone-regulated skin remodeling in *Xenopus laevis* during metamorphosis. Dev growth diff 51(4):411–427.

Tamura K, Nei M. 1993. Estimation of the number of nucleotide substitutions in the control region of mitochondrial DNA in humans and chimpanzees. Mol biol evol 10(3):512–526.

Tan MH, Au KF, Yablonovitch AL, Wills AE, Chuang J, Baker JC, Wong WH, Li JB. 2013. RNA sequencing reveals a diverse and dynamic repertoire of the *Xenopus tropicalis* transcriptome over development. Genome res 23(1):201–216.

Ter-Hovhannisyan V, Lomsadze A, Chernoff YO, Borodovsky M. 2008. Gene prediction in novel fungal genomes using an ab *initio* algorithm with unsupervised training. Genome res 18(12):1979–1990.

Thakur N, Tiwari VK, Thomassin H, Pandey RR, Kanduri M, Gondor A, Grange T, Ohlsson R, Kanduri C. 2004. An antisense RNA regulates the bidirectional silencing property of the Kcnq1 imprinting control region. Mol cell biol 24(18):7855–7862.

UniProt C. 2015. UniProt: a hub for protein information. Nucleic acids res 43:D204–212.

Vandervalk BP, Yang C, Xue Z, Raghavan K, Chu J, Mohamadi H, Jackman SD, Chiu R, Warren RL, Birol I. 2015. Konnector v2.0: pseudo-long reads from paired-end sequencing data. BMC med genomics 8 Suppl 3: S1.

Volders PJ, Verheggen K, Menschaert G, Vandepoele K, Martens L, Vandesompele J, Mestdagh P. 2015. An update on LNCipedia: a database for annotated human lncRNA sequences. Nucleic acids res 43(Database issue):D174–180.

Wang H, Iacoangeli A, Lin D, Williams K, Denman RB, Hellen CU, Tiedge H. 2005. Dendritic BC1 RNA in translational control mechanisms. J cell biol 171(5):811–821.

Wang YJ, Wang XF, Wang XZ, Li J, Wang ZS, Chen WY. 2000. High resolution late replication banding pattern of chromosomes in Rana catesbeiana. Acta Zool Sinica 46(1):115–119.

Warren RL. 2016. RAILS and Cobbler: Scaffolding and automated finishing of draft genomes using long DNA sequences. The Journal of Open Source Software. doi:10.21105/joss.00116

Warren RL, Holt RA. 2011. Targeted assembly of short sequence reads. PloS one 6(5):e19816.

Warren RL, Yang C, Vandervalk BP, Behsaz B, Lagman A, Jones SJ, Birol I. 2015a. LINKS: Scalable, alignment-free scaffolding of draft genomes with long reads. GigaScience 4:35.

Warren RL, Keeling CI, Yuen MM, Raymond A, Taylor GA, Vandervalk BP, Mohamadi H, Paulino D, Chiu R, Jackman SD et al. 2015b. Improved white spruce (*Picea glauca*) genome assemblies and annotation of large gene families of conifer terpenoid and phenolic defense metabolism. Plant j 83(2):189–212.

Weikard R, Hadlich F, Kuehn C. 2013. Identification of novel transcripts and noncoding RNAs in bovine skin by deep next generation sequencing. BMC genomics 14:789.

Willingham AT, Orth AP, Batalov S, Peters EC, Wen BG, Aza-Blanc P, Hogenesch JB, Schultz PG. 2005. A strategy for probing the function of noncoding RNAs finds a repressor of NFAT. Science 309(5740):1570–1573.

Wu TD, Watanabe CK. 2005. GMAP: a genomic mapping and alignment program for mRNA and EST sequences. Bioinformatics 21(9):1859–1875.

Yan MD, Hong CC, Lai GM, Cheng AL, Lin YW, Chuang SE. 2005. Identification and characterization of a novel gene Saf transcribed from the opposite strand of Fas. Human molec genet 14(11):1465–1474.

Zhang D, Iyer LM, Aravind L. 2011. A novel immunity system for bacterial nucleic acid degrading toxins and its recruitment in various eukaryotic and DNA viral systems. Nucleic acids res 39(11):4532–4552.

Zhang X, Wu D, Chen L, Li X, Yang J, Fan D, Dong T, Liu M, Tan P, Xu J et al. 2014. RAID: a comprehensive resource for human RNA-associated (RNA-RNA/RNA-protein) interaction. Rna 20(7):989–993.

Zhao W, Mu Y, Ma L, Wang C, Tang Z, Yang S, Zhou R, Hu X, Li MH, Li K. 2015. Systematic identification and characterization of long intergenic non-coding RNAs in fetal porcine skeletal muscle development. Sci rep 5:8957.

Zhu CB, Wang Y. 2006. A study on the chromosomes in bullfrog *Rana catesbeiana*. Acta Laser Biol Sinica 15(3): 305–307.

